# Visualization of Collagen–Mineral Arrangement using Atom Probe Tomography

**DOI:** 10.1101/2020.07.10.197673

**Authors:** Bryan E.J. Lee, Brian Langelier, Kathryn Grandfield

## Abstract

Bone is a complex, hierarchical structure comprised of two distinct phases: the organic, collagen– rich phase and the inorganic mineral–rich phase. This collagen–mineral arrangement has implications for bone function, aging, and disease. However, strategies to extract a single mineralized collagen fibril to investigate the interplay between components with sufficient resolution have been mostly confined to *in vitro* systems that only approximate the biological environment or transmission electron microscopy studies with lower spatial and chemical resolution. Therefore, there is extensive debate over the location of mineral with respect to collagen in *in vivo* mineralized tissues as visualization and quantification of the mineral in a living system is difficult or impossible. Herein, we have developed an approach to artificially extract a single mineralized collagen fibril from bone to analyze its composition and structure atom-by-atom with 3D resolution and sub-nanometer accuracy using atom probe tomography. This enables, for the first time, a method to probe fibril-level mineralization and collagen–mineral arrangement from an *in vivo* system with both the spatial and chemical precision required to comment on collagen– mineral arrangement. Using atom probe tomography, 4D (spatial + chemical) reconstructed volumes of leporine bone were generated with accuracy from correlative scanning electron microscopy. Distinct, winding collagen fibrils were identified with mineralized deposits both encapsulating and incorporated into the collagenous structures. This work demonstrates a novel fibril-level detection method that can be used to probe structural and chemical changes of bone and contribute new insights to the debate on collagen–mineral arrangement in mineralized tissues such as bones, and teeth.

## Introduction

Bone is an extraordinary material that provides vertebrates with skeletal support and essential metabolic functions, acting as a reservoir for ions involved in muscle activation to nerve signaling, and as a vessel for the production of blood cells [1]. By volume, bone can be divided into organic and mineral phases, 35% and 65 vol%, respectively. The majority of the organic phase is Type I collagen, while the inorganic phase is widely accepted to contain hydroxyapatite (Ca_10_(PO_4_)_6_(OH)_2_) with carbonate substitution. These two phases work in concert to create a hierarchical structure which is templated by the organic phase built from individual amino acids to collagen molecules to collagen fibrils and collagen fibres[2]–[4].

The mineralized collagen fibril is considered the building block for all higher-order structures within bone. However, identifying where the mineral is located with respect to the collagen fibril and what exactly that mineral is, are challenges that have plagued bone researchers for decades [5]. This is primarily due to the technical limitations associated with visualizing the mineralized collagen fibril with both sufficient spatial and chemical resolution in three-dimensions (3D). Previous work has employed high-resolution microscopy techniques, such as electron tomography, to facilitate visualization of of collagen-mineral arrangement [5]–[9]. However, these studies have been limited by the ‘missing wedge’, the inability to fully rotate a sample within the TEM, leading to distortions in reconstructions[10].Moreover, previous studies with electron tomographyhave visualized the 3D structure of bone mineral [4], [11]–[14] but not simultaneously probed for chemical information. This may account for the discrepancies seen in models for bone ultrastructure. Furthermore, there have been numerous attempts to mimic *in vivo* mineralization using *in vitro*[15]–[17], cryogenic[18], [19], and/or *in situ* approaches[20], [21], however, they are all limited by being *in vitro* systems and thus not truly equivalent to the biological *in vivo* environment.

Herein, we overcome the challenge of visualizing an individual mineralized collagen fibril from an *in vivo* system by utilizing atom probe tomography (APT) to extract and virtually reconstruct the exact chemistry and 3D structure of the collagen fibril. As shown in Fig. 1, this method enables us to study collagen–mineral arrangement from a precise site within the femur (Fig. 1A) and lamellar layers of interest (Fig. 1B), resulting in a 3D dataset displaying clear and winding collagen fibrils (Fig. 1D) that can then be explored atom-by-atom. In this paper we present the competing theories on collagen mineralization, an overview of the technical limitations associated with existing *in vitro* and 3D visualization approaches, and our findings that present new insights for collagen–mineral arrangement. Our work demonstrates that APT is a powerful technique for probing collagen mineralized *in vivo*, helping answer the questions of: (i) Where is the mineral with respect to collagen?, and (ii) What is the bone mineral composition, how does it vary spatially?

**Figure 1.**
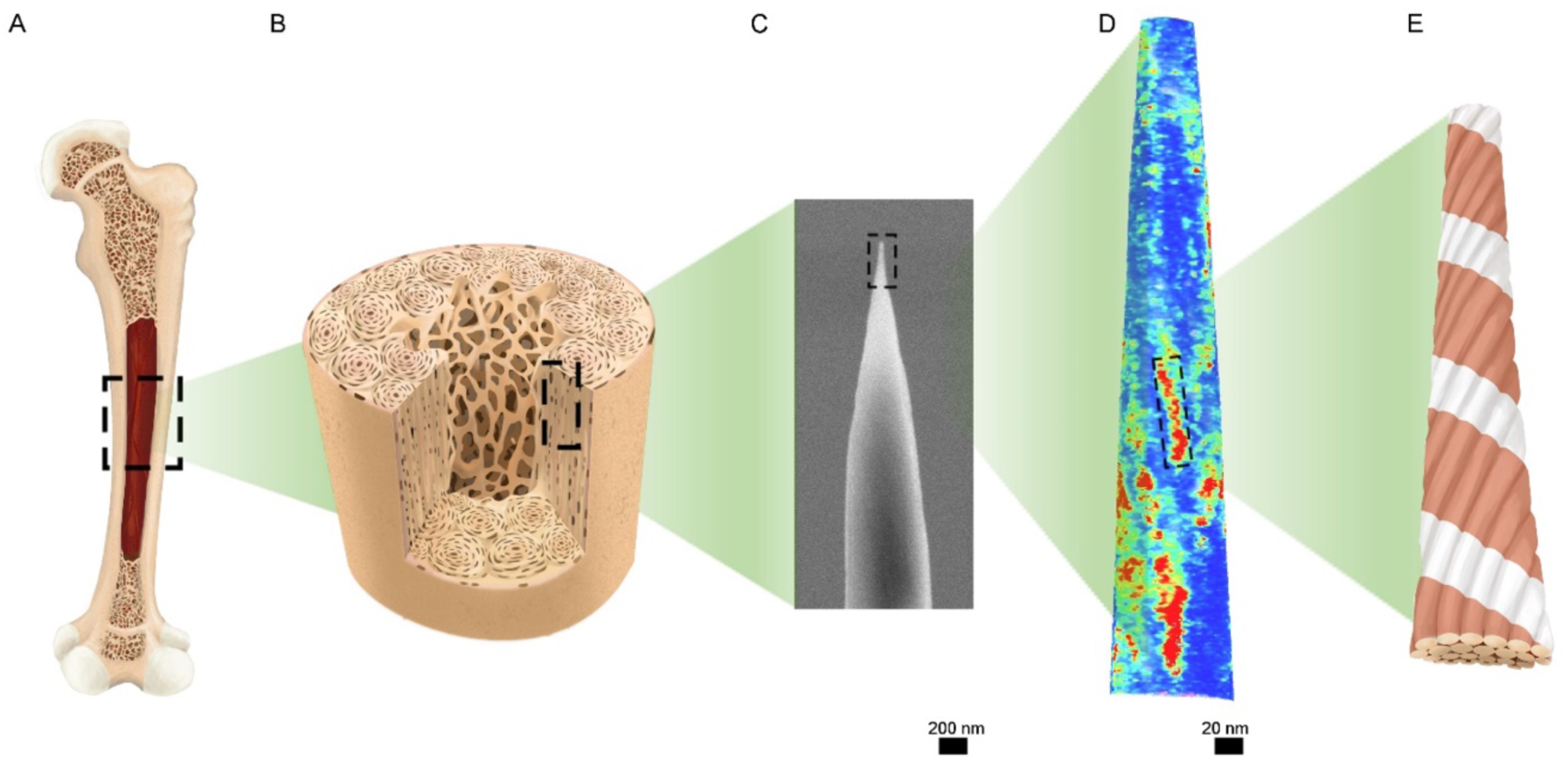
The hierarchical organization of bone as prepared for atom probe tomography. Bone is a hierarchical structure that spans the macro-to micro-to nanoscale. The femur (A) is colloquially considered to be a long bone, but upon closer inspection is comprised of complex structures, known as osteons, which act as the macroscale building blocks of bone (B). These building blocks are oriented parallel to the long axis of the femur, and within each osteon, the majority of collagen is also parallel to the femur long axis. Bone can be sectioned further using high-resolution microscopy techniques, for example by focused ion beam microscopy, a region only 50 nm in diameter with collagen oriented along its long axis is prepared (C). Using APT, collagen fibrils on the nanoscale can be visualized in bright red (D) which represent bundles of collagen molecules with their characteristic banding pattern and a right-handed helical structure (E).

### Using APT to Extract Mineralized Collagen Fibrils

Atom probe tomography (APT) is a field evaporation technique that generates 4D (3D spatial + chemical) reconstructed datasets from the ionized atoms of the sample itself, as opposed to 2D projections like electron tomography [22], [23]. Using field evaporation, atoms from a specimen surface are ejected and collected by a position-sensitive detector, giving rise to *x,y* coordinates, while simultaneously identified through time-of-flight mass spectrometry, giving rise to the mass-to-charge (*m/z*) ratio[22]. Historically, APT was not utilized for biological materials due to the requirement for a pulsed electric field to generate field evaporation which limited analysis to conductive materials [24]. However, since the advent of laser-assisted APT the technique has gained traction for analyzing various biominerals [24], [25]. APT of enamel [26], dentin [27], [28] and human bone [29]has facilitated localization of mineralization modulating elements, such as Na^+^ and Mg^2+^, within their structures or at interfaces. Human bone examined with APT showed regions of high calcium and high carbon which may correspond to the locations of mineral and collagen respectfully, but the samples were taken from the maxilla, a more disorganized type of bone, and thus the lack of *a priori* information on collagen orientation prevented a thorough analysis [29].

## Results

In long bones, such as the femur, collagen fibrils are aligned parallel to the long axis of the bone providing optimal structural support[30]. In this study, leporine bone from the femur was sectioned and using a focused ion beam (FIB), was prepared such that collagen fibrils were maintained parallel to the APT tip long axis (Fig. S1).

In this way, APT was performed and enabled long winding collagen fibrils mineralized *in vivo* to be extracted from seven APT tips, with three of the largest representing over 40 M ions each, shown here (Fig. 2A,B,C).Iso-concentration surfaces (isosurfaces) of red, yellow, and blue represent three different reconstructed surfaces high in carbon concentration, representing collagen fibrils with a measured average diameter of 20 nm that is comparable to known dimensions [31]. The chemical sensitivity of APT means that a fibril can be further isolated, where a select collagen fibril is shown without (Fig. 2D) and with (Fig.2E) corresponding calcium isosurface rendering, which represents bone mineral. Surface renderings of each of the three tips were visualized in 3D with calcium isosurfaces demonstrating the close spatial relationship throughout the entirety of the APT specimen between the collagen and mineral (Fig S2,3).On the molecular level, it is known that bundles of collagen molecules formed right-handed alpha helix structures, demonstrated by Orgel et al. that collagen molecules further formed right-handed microfibrils [32]. By using APT to extract carbon-rich collagen fibrils, our work shows that this same, right-handedness is maintained at higher ultrastructural fibril levels. Unlike previous models relying on *in vitro*, cryogenic, or *in situ* TEM, these collagen fibrils were formed and mineralized *in vivo* and their chemistry and spatial structure is simultaneously shown by APT.

**Figure 2.**
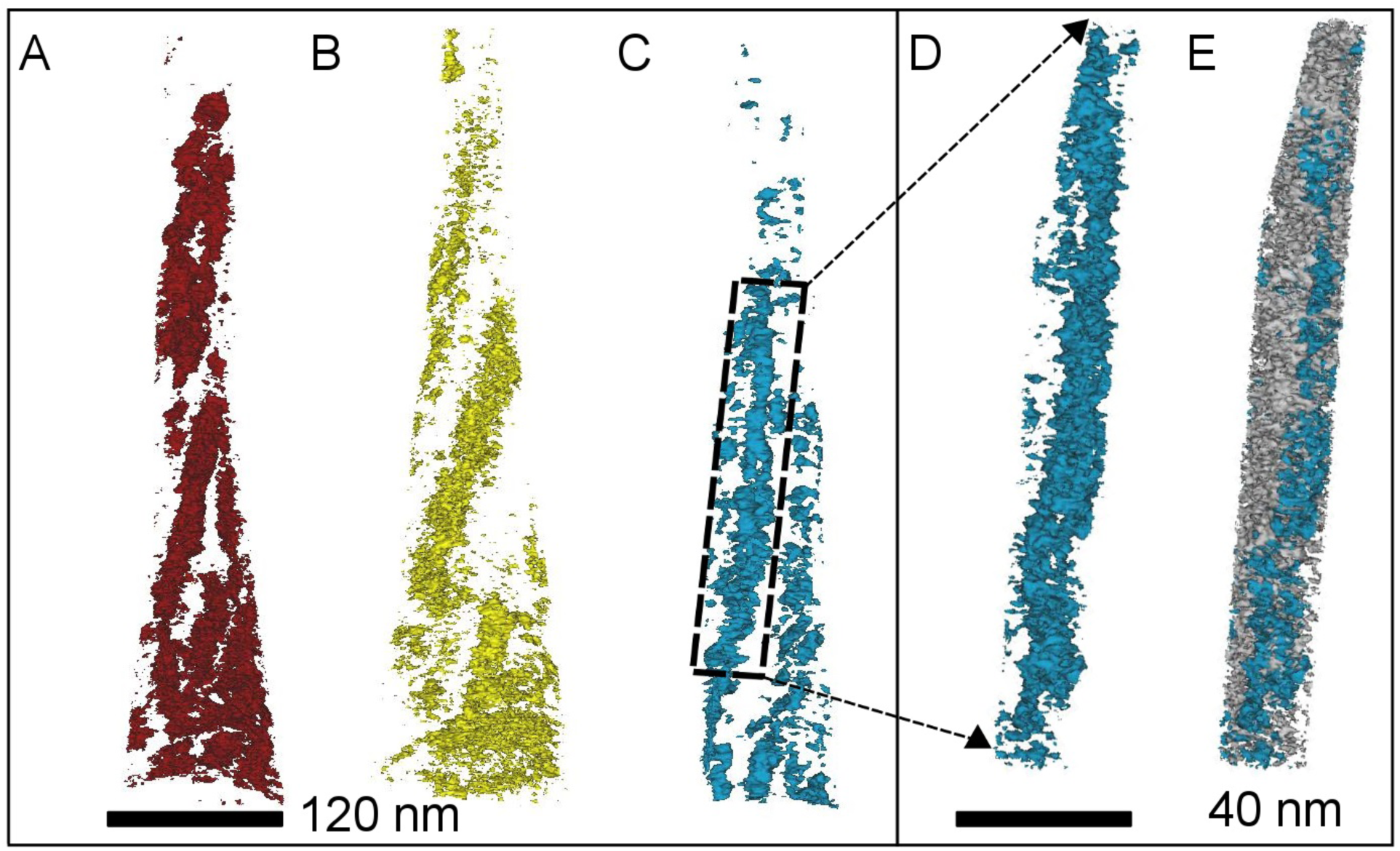
The winding collagen fibrils present with the mineralized matrix of leporine bone represented in three different reconstructed datasets. The carbon isosurfaces of three different samples (A,B,C; red, yellow, blue) (11 at%) show a fibril-like structure that is not directly parallel to the lift-out but instead appears to have some helical curvature. An individual fibril section from the reconstruction in C is isolated in D and shown along with calcium isosurface (silver; 29 at%) in E suggesting that the calcium envelopes the carbon fibril.

### Where is the mineral?

The extracted collagen fibril can be probed deeper to explore local changes in chemical composition to uncover the relationship between collagen and mineral. There have been numerous theories regarding the structure of bone, particularly how the collagen fibrils are mineralized. The dominant theories on collagen mineralization have been generated using electron tomography techniques[4], [11], [14], and broadly classify mineral placement as occurring inside the collagen fibril, intrafibrillar mineralization, or exterior to the fibril, extrafibrillar mineralization.

The chemical sensitivity and spatial accuracy of APT enables us to probe the composition in both these locations in collagen. A carbon isosurface, representing the collagen fibril (Fig 3B) is shown along with corresponding density maps of detected calcium atoms, representing the mineral (Fig. 3C), from each of the tips shown previously in Fig. 2. Corresponding density maps for carbon match the implied spatial coordinates from the isosurfaces (Fig. S4). The collagen fibril and the mineral are predominantly present in different areas of the sample (Fig. 3B,C). In general, the spaces between the carbon isosurfaces (Fig. 3B) are filled by calcium (and phosphorous, not shown) and therefore, mineral dense regions (Fig. 3C).This supports the argument of extrafibrillar mineralization dominating collagen mineralization. As in the proposed model by Schwarcz et al. the mineral surrounds the collagen in curved sheaths known as ‘mineral lamellae’ thus the crystalline mineral is completely extrafibrillar [6], [14].However, there are regions of high calcium density, most visible in the yellow specimen in Fig. 3, that correspond to the same spatial coordinates as the collagen fibril. This observation provides some evidence of intrafibrillar mineralization, and is supported by 3D visualization of the extended fibrils (Fig. S5) which show variations in concentration of calcium and phosphorus relative to the collagen fibril and some coincidence of calcium/phosphorous with collagen and nitrogen.

**Figure 3.**
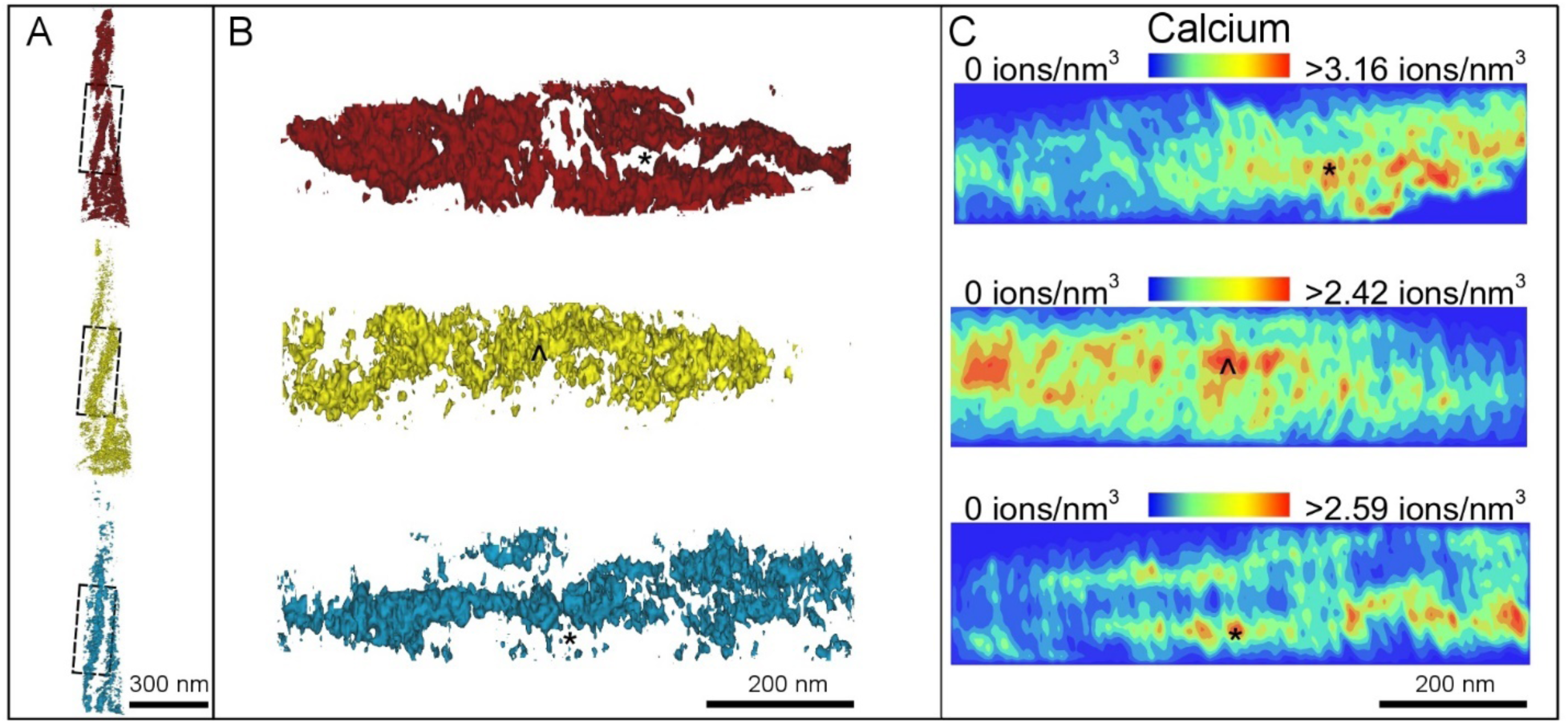
Isolated collagen fibrils (B) from three different reconstructed datasets (red, yellow, blue) (A) with corresponding calcium atomic density maps for the same region (C). A represents the original reconstruction previously shown in Figure 2, B represents the extracted collagen fibril from indicated regions in A, and C shows as-measured calcium atomic density maps. The extracted collagen fibril isosurfaces (11 at%) from all three tips show a winding structure (B). Calcium is most dense in the regions outside of the fibril (C), and example shown by the asterisk (*) on the red/top specimen. However, calcium is also present in varying high density regions within the fibrils, most visible in the yellow/middle tip, marked by arrows (^). The dimensions for B and C are identical.

The collagenous phase of bone can be broadly described as carbon and nitrogen-based, while the mineral phase is considered calcium and phosphorus-based. Atomic density maps of a representative fibril are shown for each of these four major elements through the *y,z* and *x,y* planes of the sample (Fig. 4). Density maps show a projection of the atoms in the extracted fibril. The collagen fibril is characterized by a distinct periodicity of 67 nm with gap (27 nm) and overlap (40 nm) regions between the collagen molecules which was first observed in the electron microscope and proposed by Hodge and Petruska [33]. The banding associated with collagen and its helical nature is readily visible by the non-uniformity in the density map of carbon along the *y,z* plane (Fig. 4A). Furthermore, distinct spacings between the regions of high density in the calcium (Fig. 4B) and phosphorus (Fig. 4D) maps are approximately 30 nm apart, which could suggest that gap and overlap zones identified by Hodge and Petruska are being visualized with this imaging method. Nitrogen, a major structural element in the amino acid backbone of collagen, is directly correlated (Fig. 4C) to locations of dense carbon which confirms that these long winding structures are indeed collagen fibrils. The atomic densities for both calcium and phosphorus are higher than the densities of carbon or nitrogen. Additionally, while the density of calcium at any given location is typically greater than the phosphorus density, the densest regions for both elements are highly correlated. This illustrates that although there are observable regions of variable density for calcium and phosphorus, the mineral is highly concentrated throughout the sample. When considering the calcium map, most high-density regions appear complementary to that of carbon, and along the exterior of this fibril. An example of this is shown by the asterisk marked regions in Fig 4. A,B which highlight that the central collagen fibril is flanked by dense regions of calcium and phosphorus. This is corroborated by the phosphorus map (Fig. 4D) where dense areas of phosphorus match dense areas of calcium, together providing strong evidence for extrafibrillar mineralization. Visualization of the section through the *x* and *y* plane, shown on the right of Fig. 4, shows that most of the calcium and phosphorus is on the periphery while the carbon and nitrogen are most concentrated in the centre. It is important to note that density maps are based on a projection of atoms, and therefore the *x,y* plane (on the right of the figure) represents projections along a greater thickness (the full *z* length of the sample ∼400-500nm) and therefore less correlative between Ca and P is noted. Density maps for the main elements and combinations of elements to highlight correlation are shown in more detail in Figure S7. The indication of dominant extrafibrillar mineral is in agreement with previous work that states a majority of the mineral is external to the fibril [34]. However, equally important to note, is that many of the gaps observed in the carbon map appear as repeated interdigitations of calcium, marked by arrows (Fig. 4A), which suggests intrafibrillar mineralization. Building from the work of Hodge and Petruska, Katz and Li postulated that mineral apatite crystals were arranged within the periodic collagen gap zones [34]. The most prominent theory of intrafibrillar mineralization was proposed by Landis et al. that used mineralized turkey tendon to show that inorganic mineral formed in the gap zones and nucleated outward over time[12]. Following this, Weiner and Traub demonstrated that the mineral was present in the plate-like structures that were associated with the gap zone region [35]. These theories on bone mineralization have identified mineral as being present predominantly in the gap zone regions of collagen[34], [36]–[39]. While the resolution of our datasets was insufficient to observe distinct mineral plates, the observed chemical periodicity and interdigitation of calcium and phosphorus within the collagen fibril is strong evidence of intrafibrillar mineralization within the gap zones. However, it has been noted that the total weight of mineral in bone could not be solely present in the gap zone of collagen fibrils and that some must be present externally, via extrafibrillar mineralization [34]. Therefore, these maps confirm that a combined model occurs, with both intrafibrillar and extrafibrillar mineralization occurring simultaneously *in vivo*. A periodic region with lower density of mineral measuring ∼30 nm is shown by dashed lines in Fig. 4B – this may represent the overlap zone of collagen fibrils, yet difficult to judge from the limited field of view of this sample. Correlative STEM imaging and APT, as well as collection of a longer dataset be applied to confirm this in the future.

**Figure 4.**
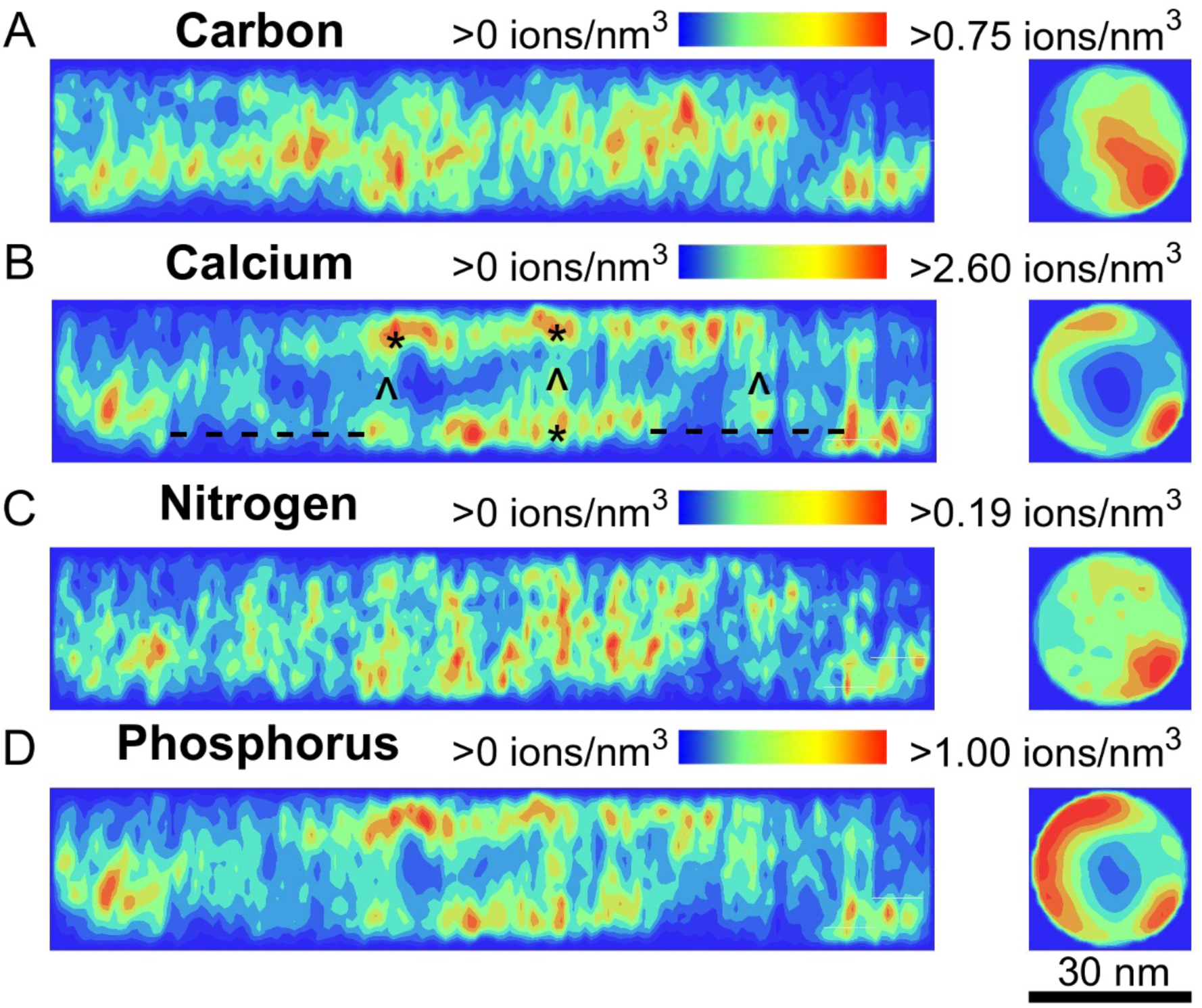
Ionic density maps visualized from the *y,z* and *x,y* planes of a collagen fibril showing distinct phases of mineral and collagen. Carbon (A) and nitrogen (C) are co-localized to form the collagen fibril while calcium (B) and phosphorus (D) are co-localized to form the mineral. The mineral is predominantly external to the collagen fibril, marked by asterisks (*) in (A) and (B). However, periodic increases in mineral density are visible within the collagen fibril, marked by arrows (^) in (A) and (B). A periodic spacing that approximates to 30 nm, indicated by dashed lines (- - -), can be observed in calcium and phosphorus maps. This may represent the overlap zone with its anticipated lower mineral content.

In order to better quantify the presence of intrafibrillar mineralization, the same sectioned fibril described above (from Fig. 4) is shown top-down from the *x,y* plane with both carbon and calcium isosurfaces (Fig. 5). At this point, it is important for us to discuss an obvious geometrical observation. The collagen fibril in cross-section, analyzed in Fig. 4 and 5, is roughly 30 nm in diameter. The range of diameter variability in collagen is actually much larger than most report, with clear evidence that fibrils range from 10-300 nm in diameter, which is highly dependent on the maturity of the collagen analyzed and the preparation method, with up to 27% shrinkage noted in some plastic embedded fibrils [40]. Diameters as small as 30 nm, similar to shown here, have been noted by second harmonic generation of rat-tail collagen [41]. It is therefore not surprising that we report smaller diameter fibrils in this work. In this cross-sectional collagen fibril view, a noticeable void in the calcium isosurface (grey) is visible when the carbon isosurface is not rendered (Fig. 5A). Whereas, rendering of the two isosurfaces (Fig. 5B) simultaneously gives an appearance similar to the ‘lacy’ mineralized crystal structure that has been frequently described [6]. The difference is that with APT, chemical sensitivity is available. Using a proximity histogram (proxigram) (Fig. 5C), the atomic concentration is measured moving into and away from the carbon isosurface (Fig. 5D; purple and red lines, respectively). The concentration of carbon greatly decreases while moving away from the collagen fibril while the calcium concentration increases, suggesting more of the calcium-rich phase is located exterior to the fibril, in extrafibrillar mineralization. In the opposite direction, moving into the collagen fibril, the concentration of carbon increases significantly while calcium drops. However, notably, the concentration of calcium and phosphorous does not completely drop off and hovers around 10 at%, indicating that there is indeed biomineral present within the collagen fibril structure, evidence of intrafibrillar mineralization. Studies by Reznikov et al. created the most recent model of mineralization which identifies mineral needles that are both in- and outside the fibril while also highlighting that differences in the ordered and disordered regions of bone can lead to alternative theories, especially when considering the small volumes that are typically analyzed in the microscope [4], [8], [37]. The results of our work show that both intrafibrillar and extrafibrillar mineralization are present in all observed APT specimens. However, recent work does indicate that collagen mineralization is highly dependent on the anatomical location even at the sub-micrometer scale. Work with plasma focused ion beam of human bone has noted mineralized clusters that span several collagen fibrils [42], and conventional FIB-SEM tomography has identified ordered and disordered regions of bone [4], which may indeed present different levels of collagen-level mineralization. Herein, we demonstrate the significant potential for APT to be extended to analysis of focused areas of bone structure, while showing for the first-time its potential to track the helical collagen fibril which high spatial and chemical resolution. Clearly further application of this technique to broader bone samples, both anatomical sites and species, will be interesting.

**Figure 5.**
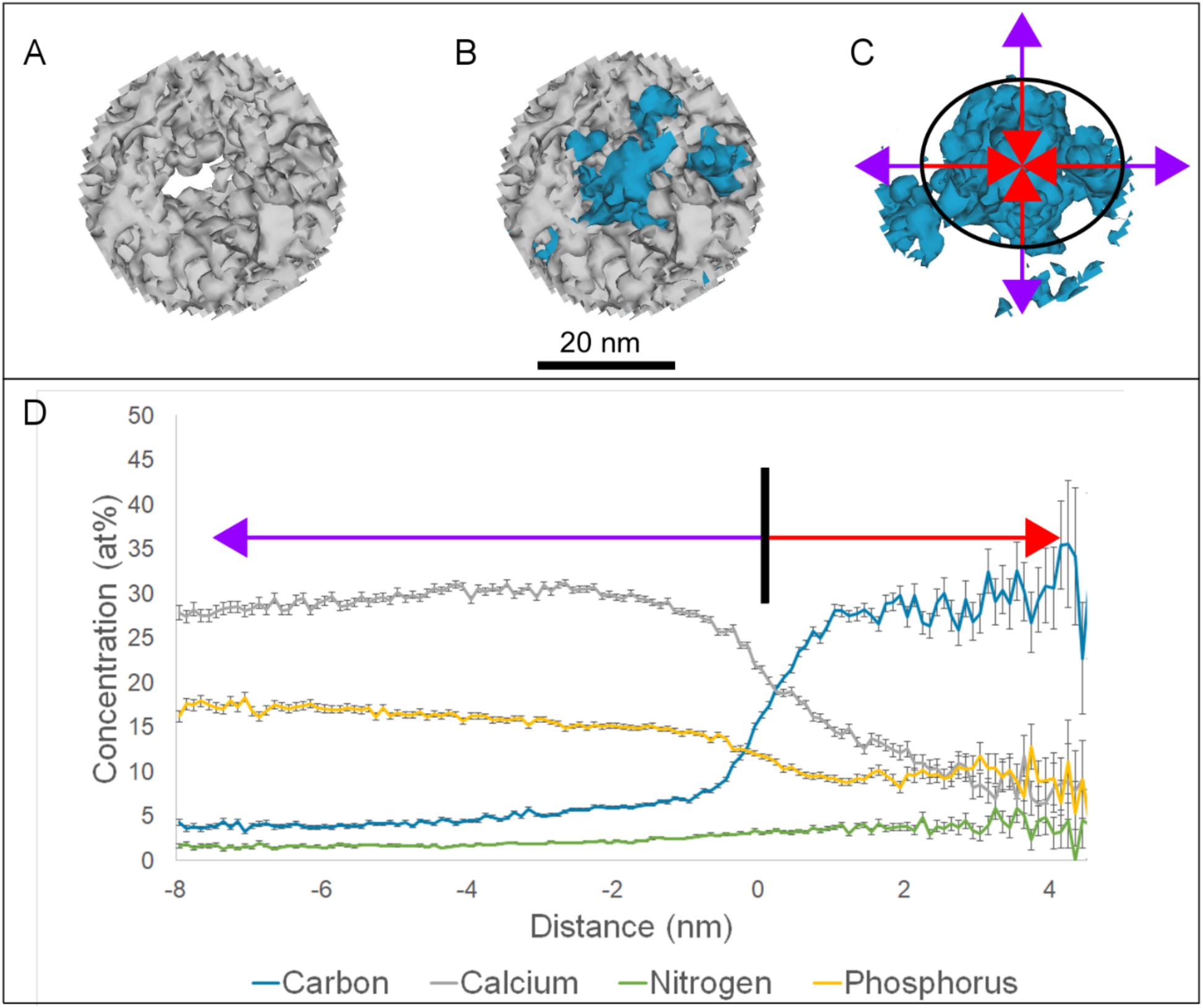
Demonstration of the spatial relationship between mineral and the collagen fibril. (A) Calcium isosurface (grey, 30.3 at%) alone showing a distinct gap in the centre of the sample. (B) Calcium isosurface rendered with carbon isosurface (blue, 11 at%) showing that the collagen fibril is enveloped completely by the mineral. (C) Carbon isosurface alone with red and purple arrows indicating the directions for the proxigram (D). Movement along the red arrow indicates measurements taken from the edge of the carbon isosurface towards the centre of the collagen fibril, while movement along the purple arrows indicates measurements taken from the edge of the carbon isosurface outwards into the extrafibrillar region. (D) Proxigram shows that while the collagen fibril is carbon and nitrogen dominant (under the red arrow), there are still significant amounts of calcium and phosphorus present suggesting intrafibrillar mineralization.

### What is the mineral?

The specific composition of the mineral phase in bone is extensively debated as is its localization. Bone apatite can be generally considered carbonate-substituted hydroxyapatite, but it is has been proposed that other phases such as amorphous calcium phosphate and octacalcium phosphate are also present in bone structure[4], [9], [34]. Amorphous calcium phosphate, in particular, was originally proposed as a precursor to hydroxyapatite during mineralization via x-ray diffraction studies by Posner [43]. This was followed by other x-ray diffraction studies which have stated that this is not ACP and may instead be poorly crystalline hydroxyapatite [44]. More recent work by Sommerdijk et al. has shown that ACP infiltrates collagen fibrils *in vitro*[18]. NMR studies have also observed ACP forming via both intra- and extrafibrillar mineralization pathways *in vitro* [45], [46].

The wealth of information provided by the APT mass spectra allows for quantification of elemental ratios. Ideally, field evaporation in an APT experiment produces monatomic ions being ejected from the APT tip one at a time. The reality – particularly for biomineralized materials with heterogeneous phases and a variety of interatomic bonds – is that often complex, molecular ions are produced during the experiment. While high quantities of monatomic ions are present in the mass spectra for bone (e.g. C or N or Ca), there are also peaks in the spectra corresponding to PO, PO_2_, or even more complex ions such as CaO_3_ or Ca_2_P_3_O_3_in various charge states (Table S1). The calcium to phosphorus atomic ratios (Ca/P) were calculated and compared to average values in the literature from varying techniques (Table 1). The average Ca/P ratio for all of the APT tips was determined to be 1.45 ± 0.02, which is very close to the theoretical 1.5 Ca/P ratio of α & β-tricalcium phosphate or amorphous calcium phosphate (ACP). This contrasts with the established 1.67 ratio of stochiometric hydroxyapatite. However, the deviation could be explained by the substitution of carbonate groups for phosphate groups within bone, as explained elsewhere[29], [44], [47].

**Table 1:** Ca/P ratio of biominerals measured using atom probe tomography and photon absorptiometry.

|  | Leporine Bone<br>– Atom Probe<br>Tomography | Hydroxyapatite<br>– Atom Probe<br>Tomography<br>[27] | Human<br>Calculus<br>Flame<br>Photometry<br>[48] |
| --- | --- | --- | --- |
| Ca/P Ratio | 1.45 | 1.66 | 1.68 |

An APT dataset, consisting of an exportable 4D point cloud, can be sectioned into regions of interest inside and outside of the collagen fibril (Fig. 6 A,B), and the mass spectra from these regions can be extracted to probe mineral chemistry. As expected, the mass spectra show there are greater quantities of mineral associated ions, such as Ca^2+^/PO^+^/PO_2_^+^ orPO_3_^+^, in the mineral region located exterior to the fibril (Fig. 6D) as opposed to within the collagen fibril (Fig. 6C). However, the mineral within the collagen fibrils had Ca/P ratios comparable to the mineral that encapsulates it. There was greater variance in the measured Ca/P ratio inside the collagen fibrils but this may be due to examining a much smaller volume that is exposed to sampling issues, wherein some cases only 20-30 atoms were being considered in calculations.

**Figure 6.**
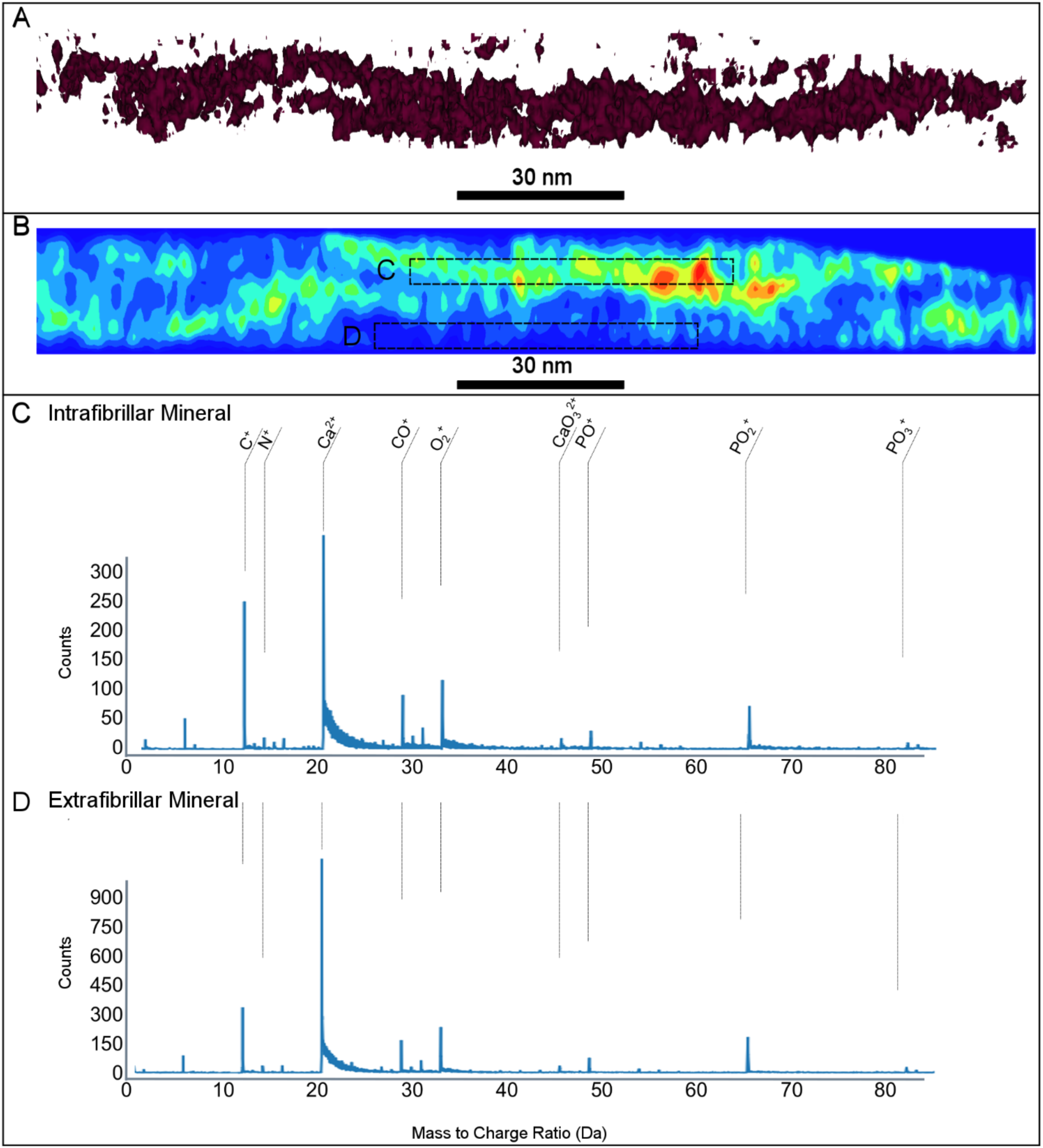
APT of a collagen fibril with representative mass spectra inside and outside of the fibril. An extracted collagen fibril (A; isosurface, 11 at%) along with carbon ionic density map (B) is shown. In the region specific mass spectra, isosurfaces extracted from the indicated ROIs, (C,D), there are greater quantities of mineral associated ions in the mineral region located exterior to the collagen fibril (D) (e.g. Ca, PO_2_) compared to that inside the fibril (C). The select ionic species indicated on the full tip mass spectra are some of those important to bone structure.

The bulk Ca/P ratios for APT reconstructed samples averaged around 1.45. However, using radial clustering analysis, groups of Ca and P atoms were selected, such that their ratios were within the ranges of either ACP (1.4-1.6) or HA (1.6-1.8) so their spatial location could be analyzed with respect to a collagen fibril. Similar analysis of proximity histograms to investigate interfaces has been conducted by Felfer and Cairney [49]. Herein, clusters labeled as ACP or HA were evaluated based on their proximity to the extracted collagen fibril by measuring the minimum cartesian distance between the two structures (Fig. 7B,C). It was observed in all datasets but one that clusters of ACP are present in greater quantities within 1.5 nm of the collagen fibril compared to HA clusters (p < 0.05). At greater distances, the proximity of the mineral to the collagen fibrils was not statistically different between ACP and HA representative ratios. This finding aligns with*in vitro* studies that note the role of ACP as a precursor to HA and the first material to form within the gap zones of collagen. However, these results also lend themselves to a discussion on the crystallinity of the mineral phase. Previous X-ray diffraction studies show that the presence of paracrystalline or poorly crystallized HA has the same Ca/P ratio as HA [44]. Our results are predicated on the assumption that the 1.4-1.6 Ca/P ratio represents ACP, but we cannot differentiate between poorly crystalline and crystalline HA using APT leaving the possibility that mineral closest to the fibril, and therefore perhaps inside the fibril, is either ACP or poorly crystalline HA. The spatial arrangement of these mineral clusters with differing stoichiometric ratios, along with their crystallinity, requires further investigation via datasets with improved mass resolution. This necessity to improve resolution is primarily due to the presence of thermal tails in APT mass spectra which have been reported with other biominerals as well [26]–[28]. In addition to clusters of mineral phases, a model for the overall winding structure of the collagen fibrils was determined by fitting the extracted bundle of collagen fibrils to a helix (Fig. 7A). The parametric equations for the bundle determined that the radius, on average, was 29.5 nm with a pitch of 16.5 nm. The model also confirmed that the helix is right-handed which agrees with known work on collagen structure [50].This model shows that APT can be used to provide a sub-nanometer and chemically accurate mathematical model parameters for the helical nature of a healthy collagen fibril. Applying this approach more broadly, APT could become an essential tool in research and study of bone diseases, particularly for specialized pathologies affecting the collagen organization, such as misfolding of the triple helix [51], Ehlers-Danlos syndrome [52], or osteogenesis imperfecta [53]. Combined with the chemical sensitivity demonstrated to probe compositional changes in Ca/P ratio, we have demonstrated APT has the power to probe both the mineral and organic framework of bone.

**Figure 7.**
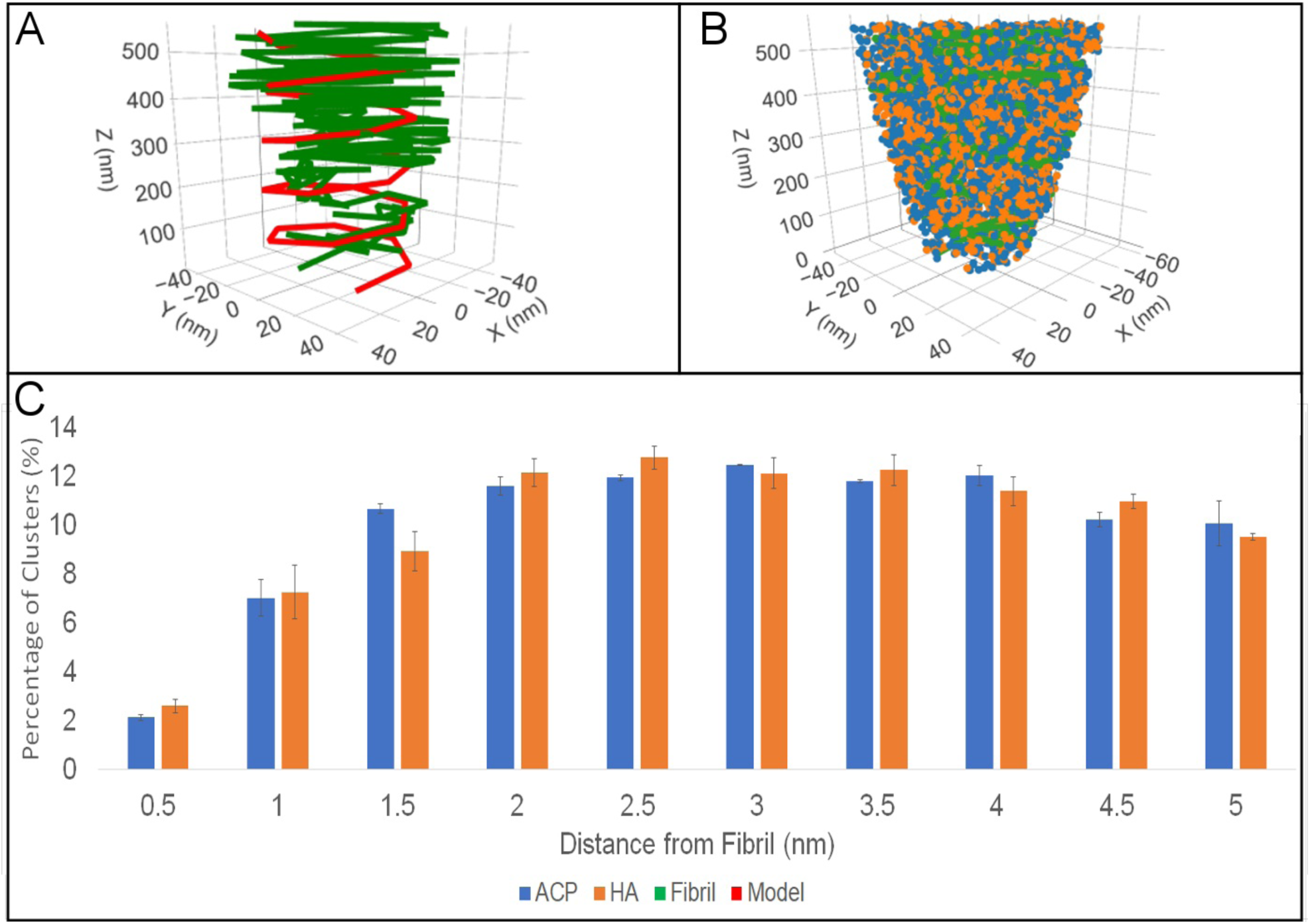
Cluster analysis of mineral proximity to collagen. Fitted helical model for a collagen fibril alongside extracted *x,y,z* coordinates of collagen fibril (A).Clusters of Ca/P with ratios of 1.4-1.6 (representing ACP) and 1.6-1.8 (representing HA) visualized using *x,y,z* coordinates (B). Histograms illustrating that up to 5 nm away from the fibril there is no difference in the amount of ACP vs. HA (C).

## Conclusion

The collagen-mineral arrangement in bone has been debated. Present theories are either based on *in vitro* systems that model but do not replicate the *in vivo* environment, or use analysis techniques without both spatial and chemical sensitivity. Herein, APT was used to clearly demonstrate the winding structure and localized chemistry of collagen fibrils mineralized *in vivo* from seven samples, each consisting of over 40 million ions. The high spatial resolution and chemical-sensitive nature of APT enabled the extraction of *in vivo* mineralized collagen fibrils from the femur of leporine bone. This work confirms with sufficient chemical and spatial resolution that both intrafibrillar and extra-fibrillar mineralization are occurring simultaneously in naturally mineralized bone, refuting models that insist only one or the other exists. Ca/P ratios were also shown to vary with respect to the collagen fibril location, suggesting possible increases in amorphous or low crystallinity apatite closer to, or perhaps within, the collagen fibril. Therefore, APT has shown the capacity provide newfound insight to the structure and chemistry of organic and inorganic materials, such as the fundamental building block of bone shown herein, the mineralized collagen fibril.

## Materials and Methods

### Preparation of Bone Samples

A single femur of a fresh-frozen wild (age-unknown) leporine bone (*Lepus americanus*), commonly known as snow-shoe hare, was received with institutional ethical approval, was cut transversely to its long axis using a slow-speed saw (Isomet LS, Buehler Ltd.) and diamond blade to generate a cross-section from the mid-diaphysis region. Bone marrow was removed from the samples by submerging them in a 3% sodium hypochlorite solution (in Milli-Q water) for 3 hours, as conducted by other seminal works highlight bone ultrastructure and therefore considered an acceptable level of chemical treatment which would not alter collagen-mineral morphology extensively in the bulk of the sample [54]. This treatment was only used on the bulk sample of whole femur, and not the exposed slices used in this work, therefore further mitigating the possibility for altering the collagen and mineral. Samples were subsequently dehydrated in a graded series of ethanol from 25-100% (in Milli-Q water) as previously described [55]. Following dehydration, the samples were infiltrated with resin (Embed 812, Electron Microscopy Sciences) over 4 days before being placed in a 50°C oven overnight. Tissue blocks were polished manually (400, 600, 800, 1200 grit SiC paper and subsequently cloth with 0.3 µm alumina suspension (MetLab)) until embedded bone was exposed. Samples were sputter-coated with 10 nm of chromium prior to imaging. Scanning electron microscopy (SEM) in backscattered mode was performed at 10 kV (JEOL 7000F) to select regions of interest. A typical ROI is outlined in Fig S1, which was selected from the endosteal surface of lamellar bone (lamellar bone visible in Fig S1 C and D). The relative age of this region is unknown, however, backscattered electron imaging does not indicate that the ROI is significantly mineral rich (older) or mineral deficient (recently formed) compared to surrounding regions. In future work, identifying regions of graded maturity would be interesting, yet very challenging, and indeed beyond the scope of the proof-of-concept work herein.

A dual-beam focused ion beam (FIB) microscope (NVision40, Zeiss) was utilized to prepare tips for APT according to previous studies [29]. An *in-situ* lift-out method (Fig. S1) was used to obtain site-specific samples. Following lift-out, samples were mounted to pre-sharpened Si coupons or W needles, and milled to create tips with appropriate sharpness for APT (diameter approximately < 150 nm). Tips were prepared such that the collagen fibrils were aligned parallel to the long-axis of the tip. Each of the seven tips presented in this work were obtained from the same lift-out.

### Atom Probe Tomography

APT was performed using a LEAP 4000XHR (Cameca, WI, USA) microscope operating in laser pulsing mode (355 nm UV, 125 kHz pulse rate, 50 pJ pulse energy) at a stage temperature of 59.6 K under ultra-high vacuum (< 10^−8^ Pa). An evaporation rate of 0.005 ions/pulse (0.5%) was maintained by controlling the static DC potential applied to the specimen, which typically ranged from 2.0-4.3 kV. A total of seven tips were successfully run, with total evaporated ions for the runs ranging from 36-50 million ions, with three of the largest tips highlighted in this work.

Reconstructions were performed using IVAS 3.6.10a (Cameca, WI, USA). Each reconstruction was completed using tip profile radius evolution, utilizing SEM images of the tips taken before and after each APT run to guide the reconstruction. An image compression factor of 1.65 was used throughout. There were a number of peaks that were not possible to range due to challenges associated with running a biological sample through the atom probe microscope, such as large thermal tails. As such, the average detector efficiency was adjusted based on the number of peaks that were successfully ranged, resulting in a modified detection efficiency parameter of ranging between 0.07 and 0.1 for each tip. For analysis, all identified ions were used, a complete list is provided in Table S1.

To ensure the accuracy of reconstructed datasets, correlative electron microscopy was performed. SEM images taken before and after APT runs were compared to determine the area that was evaporated. For some tips, electron tomography was attempted in a Titan 80–300 TEM operating in STEM mode (FEI Company, The Netherlands) at 300 keV at 2° tilts from -65 to 65 ° (Fig. S6). While the tips were suitable for runs in the APT afterwards, the electron tomography acquisition damaged the tips during its imaging making reconstruction not possible.

### Statistics and Model Generation from APT Data

Helical regression and mathematical calculations of cartesian distances between clusters of various Ca and P ions, as well as Ca:P ratios and the collagen fibrils was computed using R (version x64 3.6.1). Clusters of Ca:P were extracted from IVAS using Ca, PO, PO_2_, PO_3_ and P clustering centres. The following packages were used in the analysis: dplyr, plot3d, plot_ly, rgeos, rgl, and sp.

## Acknowledgements

This work was supported by the Natural Sciences and Engineering Research Council of Canada (NSERC) (RGPIN-2014-06053), and the Ontario Ministry of Research, Science and Innovation (Early Researcher Award ER17-13-081). Microscopy was performed at the Canadian Centre for Electron Microscopy (CCEM) at McMaster University, a facility supported by NSERC and other government agencies. We thank Dr. Andreas Korinek for assistance with STEM tomography experiments. We thank Dr. Cheryl Quenneville and Dr. Solomon Amuno for providing the bone samples. We thank Marcia Reid for sample preparation assistance.

## Author Credit Statement

Conceptualization; K.G. and B.E.J.L Methodology: B.E.J.L., B.L., K.G

Data curation; B.E.J.L., B.L.

Formal analysis; B.E.J.L., B.L.

Funding acquisition and supervision; KG

Writing: original draft; B.E.J.L., K.G.

Writing: review & editing; B.E.J.L., B.L., K.G.

## Ethics Declaration

The authors declare no competing interests.

## Supplementary Information

**Figure S1:**
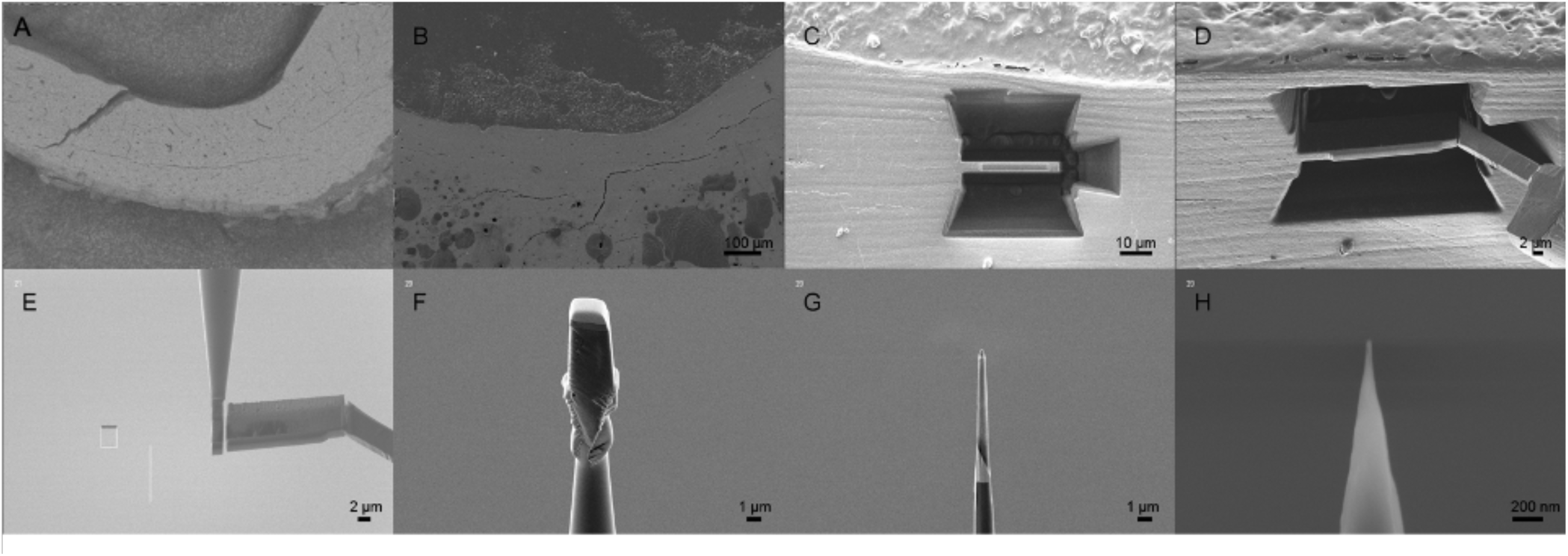
Preparation of sectioned bone into APT tip. A region of interest was identified in SEM near the edosteal surface (A/B). The area around the ROI was milled away using Ga ions (C) before being extracted using a micromanipulator (D). The lamellar nature of the ROI is clearly evident from the lamellar lines visible in C and D. The removed section (E) was split into several pieces and attached to an APT coupon (F). Each were sharpened (G/H) into the needle shape required to perform APT.

**Supplementary Table S1:**
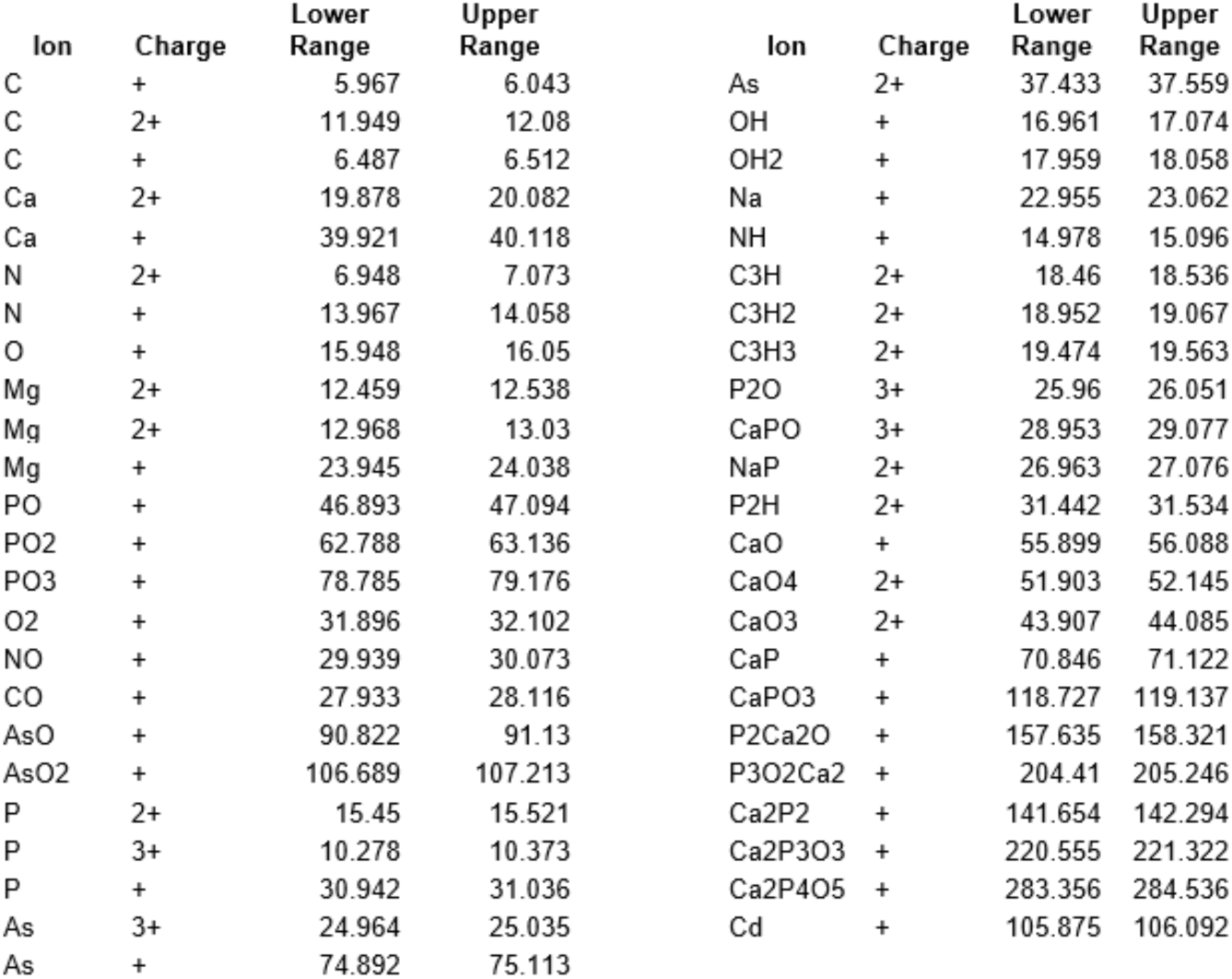
List of ranged ions.

**Figure S2:**
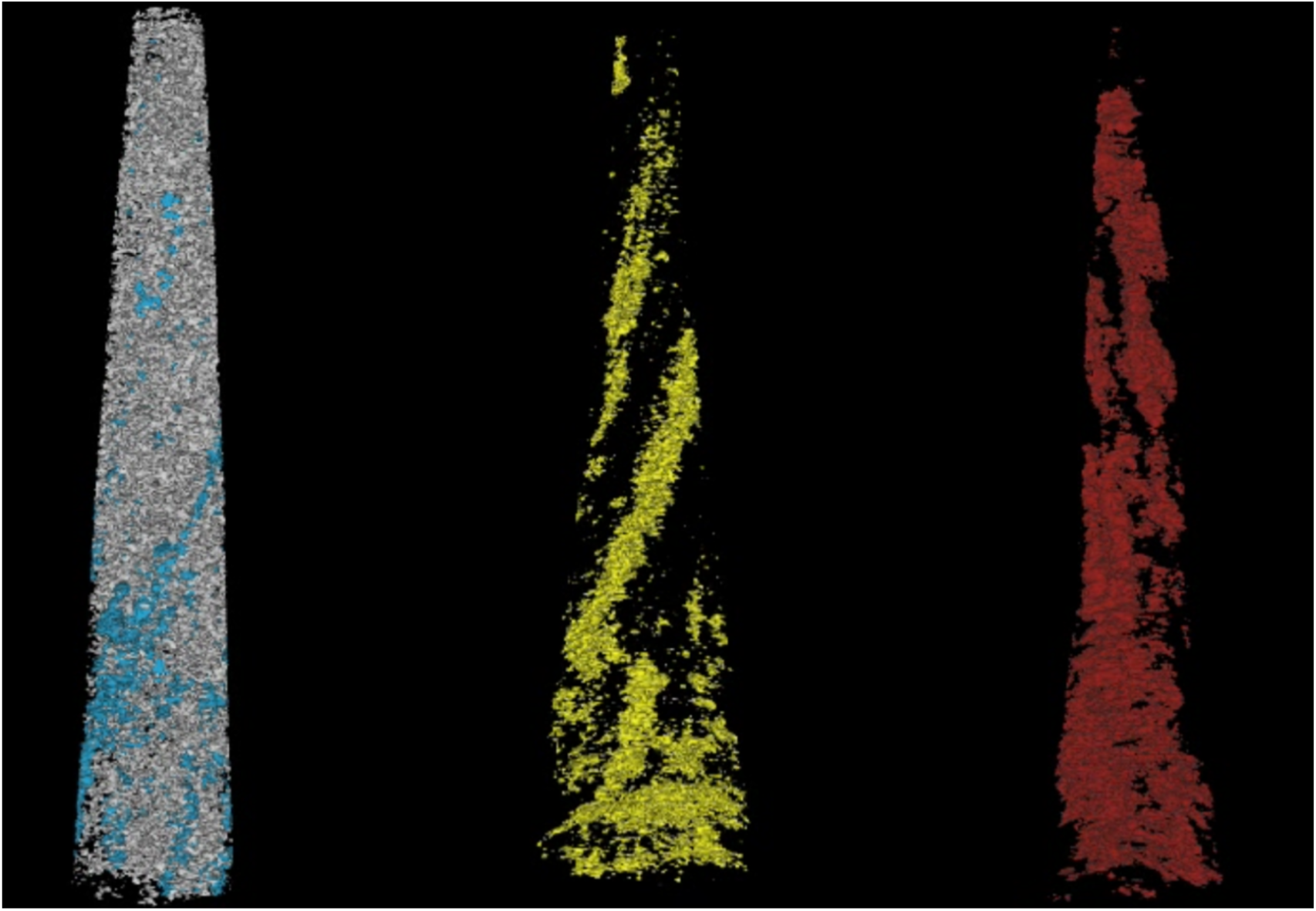
Rotated Isosurfaces Renderings of 3 APT tips: C (11 at%, blue, yellow and red) and Ca (29 at%, grey) *See Supplementary Video 1.

**Figure S3:**
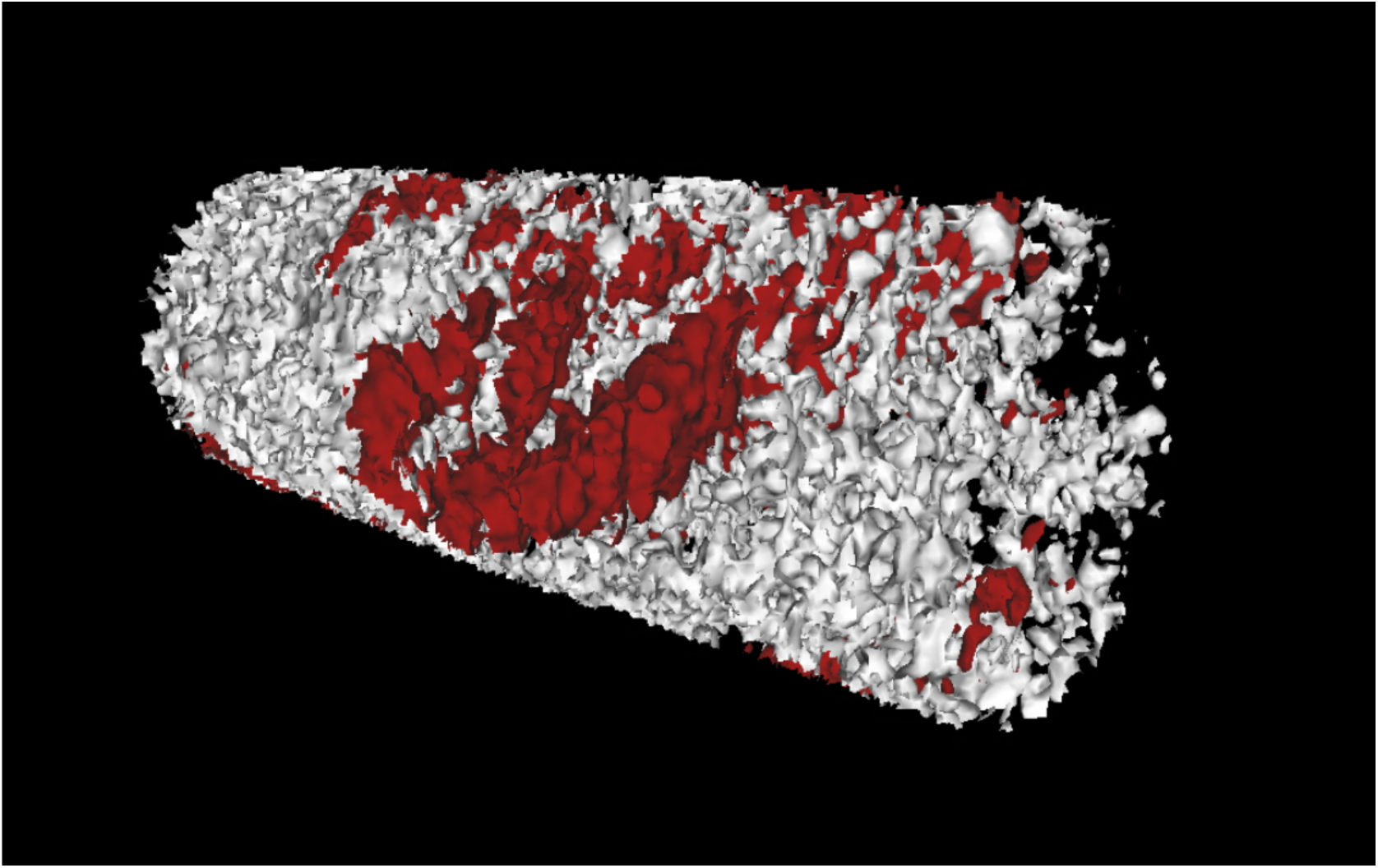
Single isolated fibril section showing C (11 at%, red) and Ca (29 at%, grey). The encapsulation of collagen, represented by carbon, is clear. *See Supplementary Video 2.

**Figure S4.**
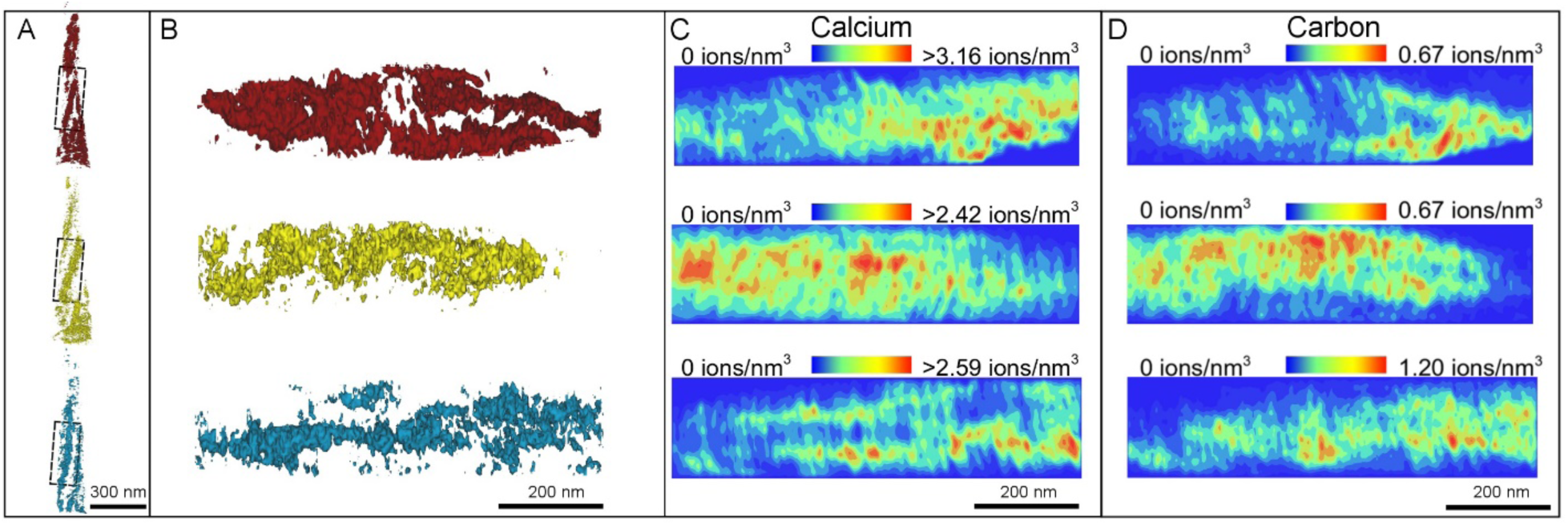
Isolated collagen fibrils from three different reconstructed datasets with corresponding calcium and carbon atomic density maps for the same region. Figure 3-3 with carbon ionic maps added. Dimensions for B,C, and D are the identical. D shows that the carbon density matches the spatial coordinates of the isolated collagen fibrils shown via isosurface rendering in B.

**Figure S5.**
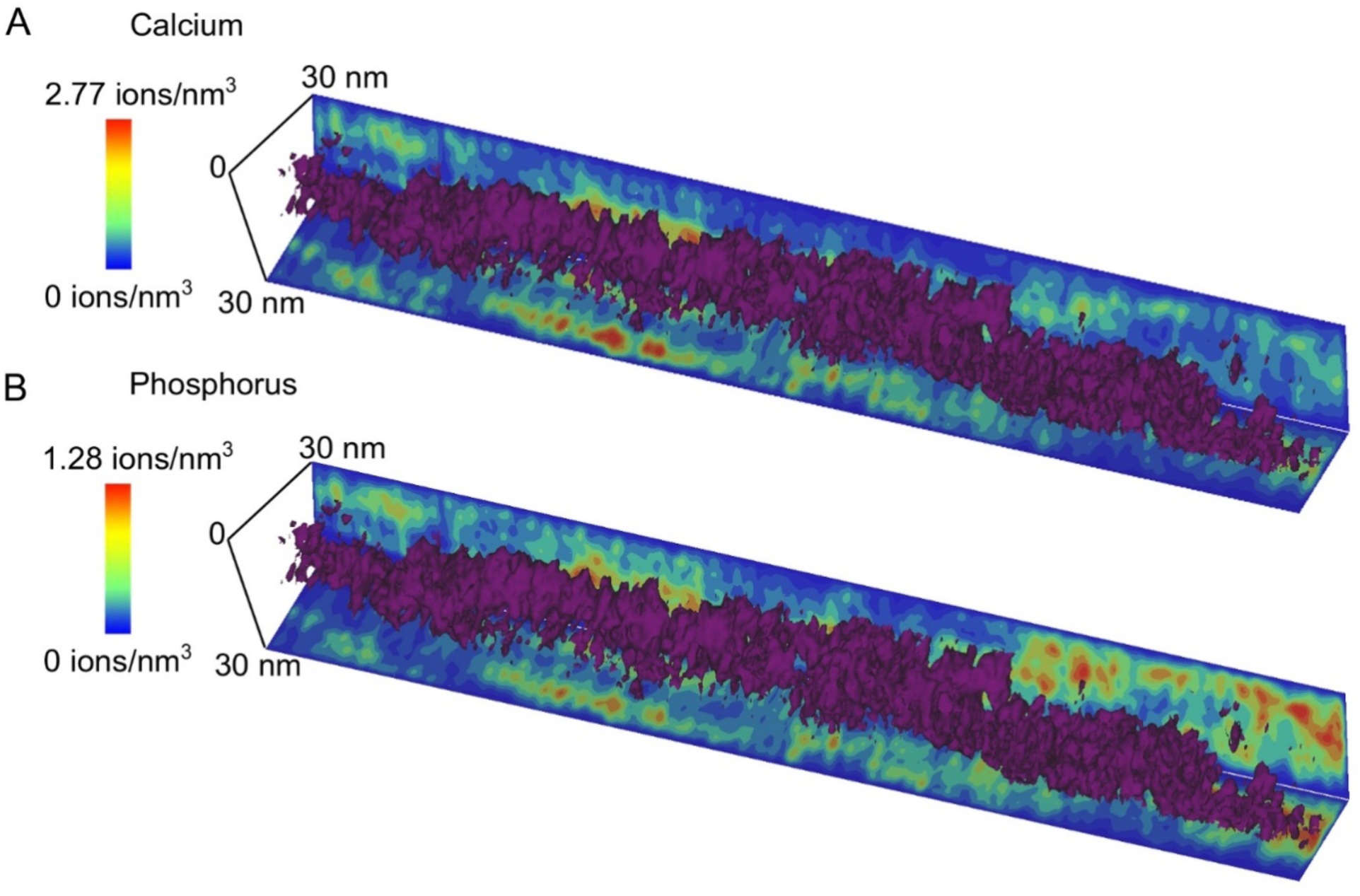
Extracted collagen fibril superimposed on atomic density maps of calcium and phosphorous. An extracted fibril, 285 nm in length, is shown to both incorporate and be encapsulated by calcium and phosphorus demonstrating that while extrafibrillar mineralization dominates, intrafibrillar mineralization is also occurring in bone.

**Figure S6.**
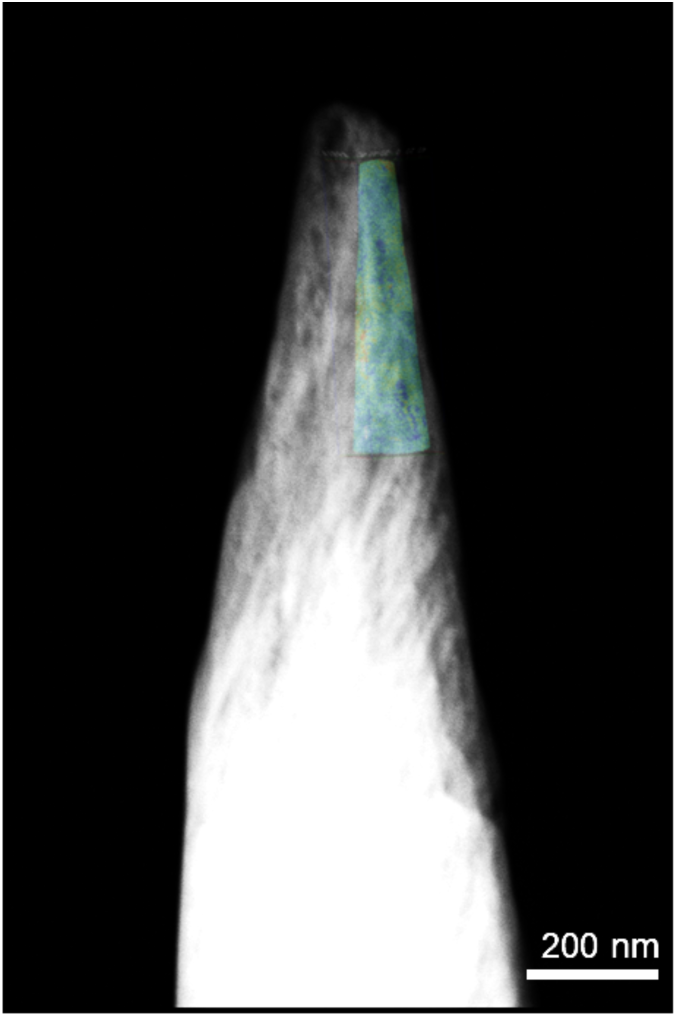
Image from STEM-HAADF electron tomography with overlaid APT volume. APT detector and evaporation efficiency reconstruction parameters were altered using structural information from STEM imaging and tomography to improve APT reconstruction accuracy. A representative APT tip is shown overlapping the STEM image in cyan.

**Figure S7.**
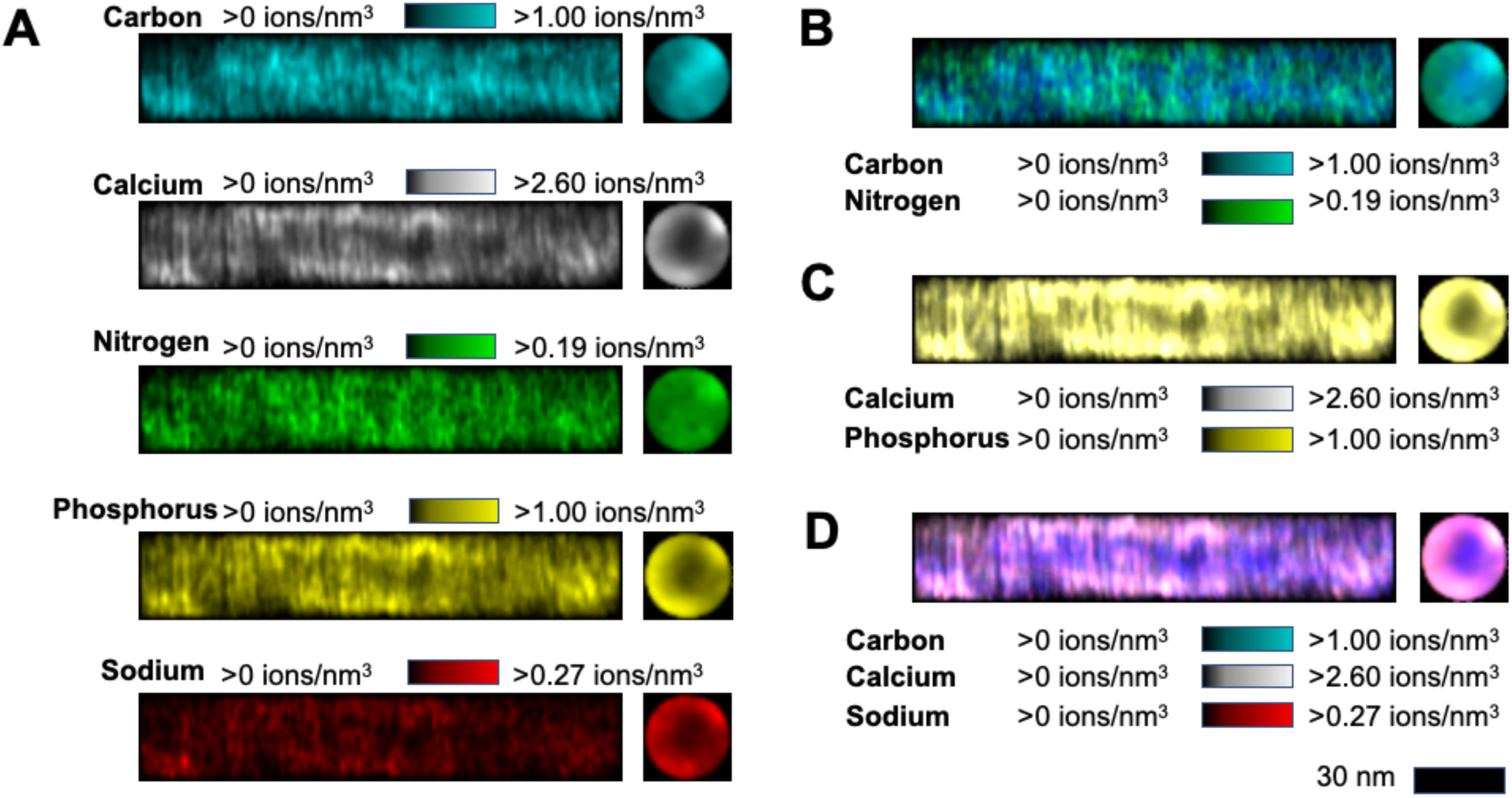
Composition mapping of elements in leporine bone. A) The distinct main elements (Carbon, Calcium, Nitrogen, Phosphorus, and Sodium) comprising the collagenous and mineral components of bone. B) Overlay of carbon and nitrogen maps showing co-localization. C) Overlay of calcium and phosphorus maps, showing co-localization. D) Overlay of carbon, calcium and sodium maps, showing sodium predominantly in the mineral phase but also in near the periphery of the collagenous zone.

